# Type IV pilus length determines virulence by regulating a hidden subpopulation of non-contributing filaments

**DOI:** 10.64898/2026.04.02.716181

**Authors:** Zil Modi, Jessie Reed, Ahmed O. Yusuf, Tyler Payne, Neel Chalemela, Sam Denison, Matthias D. Koch

## Abstract

Many clinically important bacterial pathogens, including *Pseudomonas*, *Vibrio*, *Neisseria*, and *Acinetobacter* species, employ dynamic extracellular appendages called type IV pili (T4P) to facilitate virulence through cyclical extension and retraction of pilus filaments. To dissect how T4P dynamics govern pathogenesis, we engineered a genetic system to precisely tune pilus length across a continuum. We demonstrate that pilus length critically determines four major T4P-dependent virulence traits in *Pseudomonas aeruginosa* (motility, surface sensing, biofilm formation, and phage infection) and reveal a hidden subpopulation of pili that are unable to interact with environmental substrates or host cells, rendering them non-contributing to any T4P-mediated function. Integrating molecular dynamics simulations, we show that low inner-membrane abundance of the major pilin forces the extension mechanism into transient idle states, restricting both velocity and final length. Molecularly, this finding reveals how two key biophysical parameters, pilin abundance and diffusion, impose a fundamental physical constraint on T4P assembly, and that regulating pilin abundance presents a strong lever over regulating pilus count for controlling the amount of functionally contributing filaments. Contrary to the prevailing view that retraction force generation primarily dictates T4P-mediated behaviors, our results establish extension dynamics as the overlooked bottleneck constraining all retraction-enabled virulence traits, with population heterogeneity in length enabling adaptive bet-hedging.

## Introduction

Type IV pili (T4P) are dynamic extracellular filaments that enable bacteria to interact with their environment and host tissues. These fibers assemble from a pool of major pilin subunit at a multiprotein envelope-spanning machine that allows extension from the inner membrane into the extracellular space [1–5]. Through repeated cycles of extension and retraction, T4P power key virulence behaviors, including twitching motility for surface exploration and dispersal [1, 6, 7], biofilm formation that promotes colony protection against host defenses and antibiotics [8–12], DNA uptake for horizontal gene transfer of resistance and virulence traits [13–16], and surface sensing to coordinate pathogenic responses [17–20]. Consequently, T4P-equipped pathogens are notoriously difficult to treat, as their adaptability drives rapid colonization and systemic infection.

T4P-mediated virulence traits are widespread across Gram-negative pathogens (*Pseudomonas*, *Vibrio*, *Neisseria*, *Acinetobacter*, and others) [17, 19, 21–28], but *Pseudomonas aeruginosa* stands out as a unique and powerful model system: it robustly employs four major T4P-dependent behaviors (twitching motility, surface sensing, biofilm formation, and phage infection), making it ideal for dissecting how T4P dynamics control pathogenesis [9, 19, 29–36].

In *P. aeruginosa*, the major pilin subunit PilA is polymerized to extend the pilus fiber and depolymerized during retraction. PilA expression is regulated by the two-component system PilSR. When PilA levels are low, the sensor kinase PilS auto-phosphorylates and activates the response regulator PilR, which drives *pilA* transcription in a *rpoN*-dependent manner [36–38]. This feedback is thought to maintain a constant inner-membrane pool of PilA for pilus assembly [39]. However, the functional consequences of changes in PilA abundance and how they affect T4P dynamics and virulence remain poorly understood.

The T4P assembly complex includes the PilMNOP alignment subcomplex linking the cytoplasmic platform PilC to the outer-membrane secretin PilQ [40–42]. Hexameric ATPases PilB and PilT engage PilC to drive extension and retraction, respectively [41–45]. ATP hydrolysis by these motors induces conformational changes in PilC that incorporate or remove PilA subunits from the inner membrane [2, 4, 46]. Recent studies suggest that the switch between extension and retraction is primarily driven by stochastic and mutually exclusive binding of PilB and PilT to PilC [5, 47, 48], however, mechanosensitive effects triggered by surface contact may also contribute to modulating this switch [19, 33, 49, 50].

Retraction generates a large range of forces from <10 pN to >100 pN per pilus or nN in bundles for different species, making T4P one of the most powerful known molecular motors [13, 17, 18, 51–57]. Optical tweezers studies have revealed detailed insights into the molecualr mechanisms driving retraction, including bimodal velocities, force-dependenta switching to extension, and two-dimensional tug-of-war in twitching [58, 59]. However, the flexibility of T4P fibers limits optical tweezers for extension studies, leaving the molecular processes governing pilus extension far less explored.

T4P research has primarily emphasized retraction as this is the force-generating step, with extension often viewed as a prerequisite rather than a regulatory bottleneck. Yet retraction depends on prior extension, which determines pilus length and the fraction of pili that is available for functional interactions. Our incomplete understanding of extension mechanisms thus limits insight into T4P-mediated virulence. Here, using *P. aeruginosa* as a model, we developed a genetic tool to precisely tune pilus length while visualizing dynamics in real time. This approach reveals that pilus length critically governs virulence traits, uncovers a hidden subpopulation of non-contributing short pili, and identifies diffusion-limited PilA supply as the biophysical constraint that biases extension and makes length regulation more efficient than count regulation for controlling functional output.

## Results

### Clonal populations display a broad heterogeneity of pilus activity and PilA levels

To investigate the molecular mechanisms underlying pilus extension, we quantified pilus number and length in individual cells using cysteine-maleimide-based fluorescent labeling of pili in a *pilA*-A86C background [18, 48, 60]. Consistent with prior studies, pilus count and length distributions were exponential, with a strong bias toward cells producing few and short pili [13, 48] (Figure 1A,B). A typical cell extended 1–5 pili within a 30-second observation window, with lengths ranging from 0.2 to 1.5 µm, although some cells produced up to 13 pili reaching lengths of up to 4 µm.

**Figure 1.**
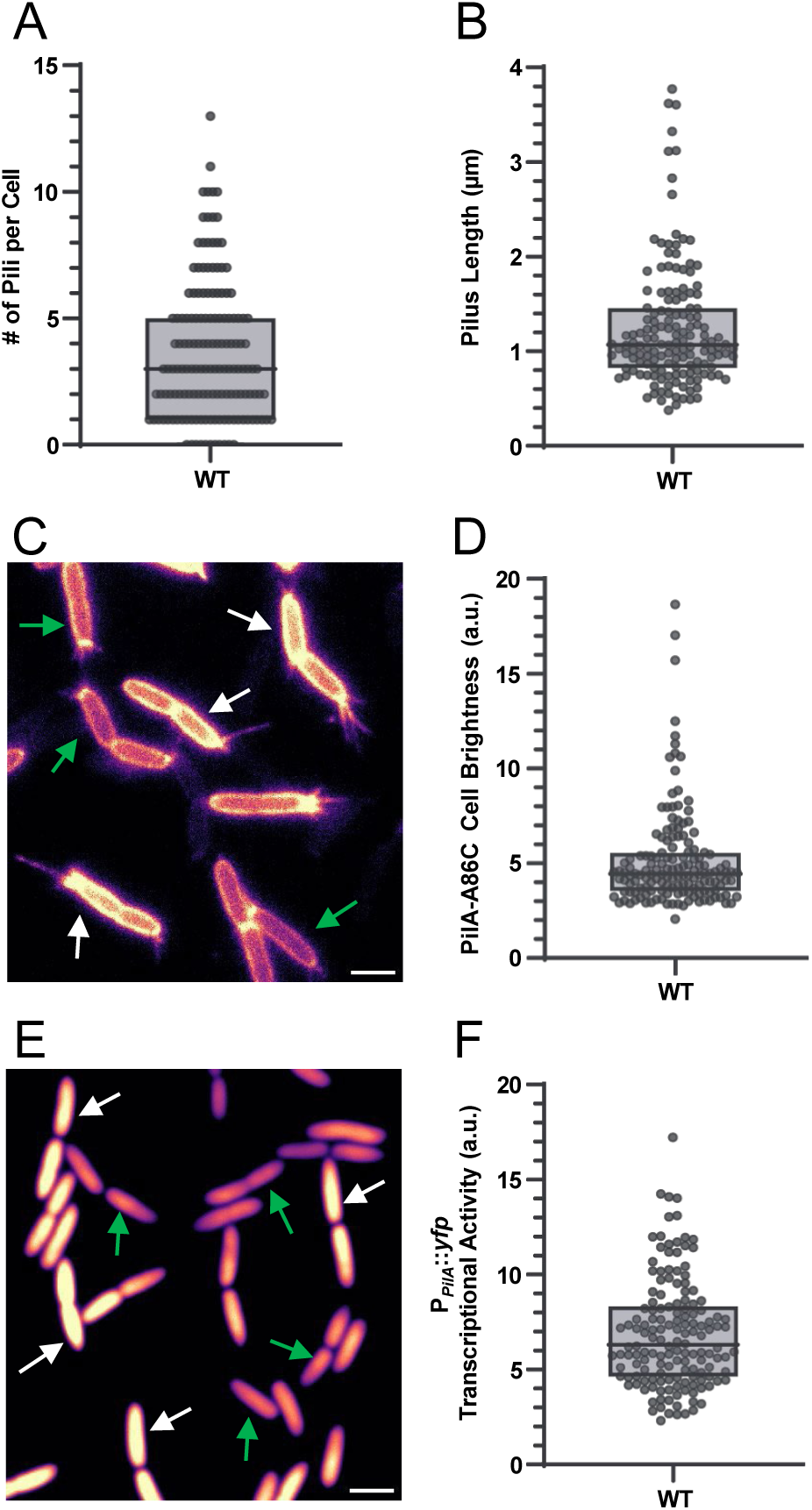
Pilus dynamics and major pilin expression display a 10-fold difference between individual cells in a clonal population. **A)** Total count of individual pili per cell, quantified from 30-second live-cell movies. **B)** The maximum extension lengths of individual pili within the same 30-second live-cell movies as in (A). **C)** Representative fluorescence still image of cells with Alexa488 labeled PilA-A86C cysteine knock-in mutant. **D)** Brightness of individual cell bodies with Alexa488 labeled PilA-A86C. **E)** Representative fluorescence still image of *pilA* transcription using a *P_pilA_::yfp* reporter in single cells. **F)** Brightness of *P_pilA_::yfp* reporter in single cells. **Box plots:** Boxes represent median and interquartile range (25th–75th percentiles) of N = 150 cells/pili from three independent biological replicates. **Images:** White arrows: bright cell bodies, indicating high PilA protein levels (C) or gene expression (E). Green arrows: dim cell bodies, indicating low PilA protein levels (C) or gene expression (E). Scale bars: 2 µm.

We also observed marked variation in cell body fluorescence: some appeared dark while others were significantly brighter (Figure 1C). As this labeling targets the major pilin PilA, cell body brightness reflects intracellular PilA levels. Quantification revealed an exponential distribution of brightness from ∼2 to 20 arbitrary units (a.u.), with most cells at 3–6 a.u. (Figure 1D). To rule out non-specific membrane labeling artifacts, we confirmed similar heterogeneity in single cells using a transcriptional reporter *P_pilA_*::*yfp*, showing *pilA* promoter activity spanning roughly one order of magnitude (Figure 1E,F). Together, these data demonstrate substantial cell-to-cell heterogeneity in PilA abundance within the population.

### Abundance of the major pilin PilA tunes pilus dynamics

The observed cell-to-cell variation in PilA abundance was unexpected, as PilA expression is thought to be tightly regulated by the PilSR two-component system, which provides feedback between protein levels and gene expression [37, 38, 61]. To directly test how PilA abundance influences pilus extension, we constructed an arabinose-inducible allele (*P_bad_*::*pilA*-A86C) integrated into the chromosome in a *pilA* background. Western blotting confirmed a tunable range that recapitulates the natural PilA heterogeneity observed in individual cells: no inducer (0% arabinose) yielded ∼10% of wild-type (WT) PilA levels, while concentrations above 0.4–0.5% plateaued at ∼130% of WT. Induction of 0.3% arabinose closely matched WT levels (Figure 2A, Supplementary Figure 1).

**Figure 2.**
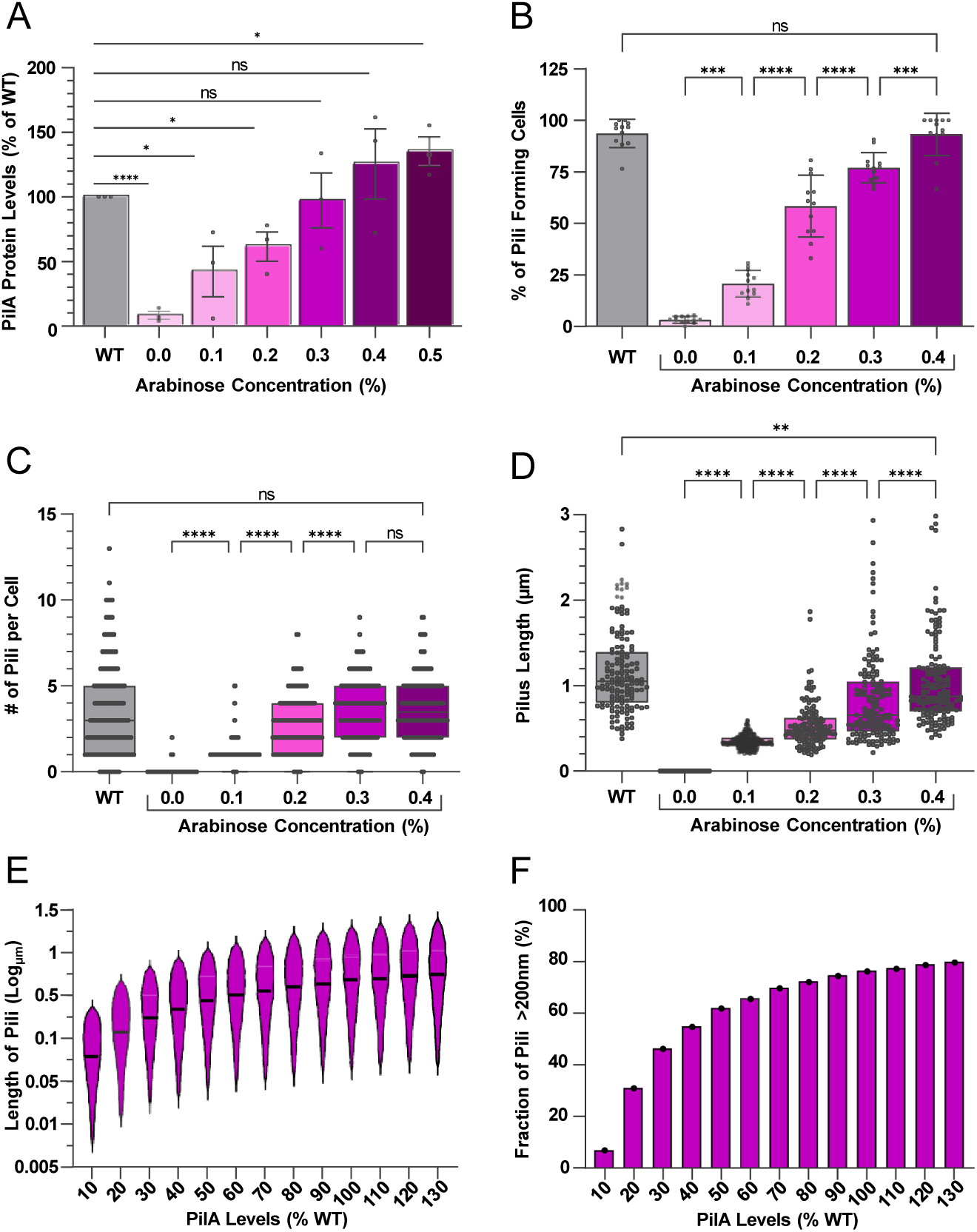
PilA protein levels determine the activity and dynamics of pili in individual cells. **A)** PilA protein titration using *P_bad_*::*pilA* in a *pilA* background determined by Western blotting using a polyclonal PilA antibody. Bars represent mean ± SEM of three biological replicates (indicated as black dots). **B)** Fraction of cells forming at least one pilus in a 30 second window as a function of PilA titration (same strain as in panel A). Bars represent mean ± SD of N = 12 technical replicates from three biological replicates. **C)** Total number of individual pili in a 30 s window in single cells as a function of PilA titration (same strain as in panel A). Boxes show median and 25th–75th percentiles of N = 150 cells from three independent biological replicates. **D)** Maximum pilus extension lengths for individual pili as a function of PilA titration (same strain as in panel A). **E)** Pilus lengths for 10,000 simulations of individual pilus extension as a function of PilA levels in the inner membrane. Violin plots show median (horizontal line) and 5% to 95% percentiles. **F)** Fraction of simulated pili that reached a maximum length of >200 nm from the 10,000 simulated pili in E as a function of PilA levels. **A)** Statistical tests were performed using unpaired t-test. **B) – D)** Statistical tests were performed using one-way ANOVA. Significance: (ns) not significant, P > 0.05; (*) P < 0.05; (**) P < 0.01; (***) P < 0.001; (****) P < 0.0001.

We next examined the impact of PilA levels on pilus dynamics. The fraction of piliated cells increased with PilA abundance (Figure 2B): at 0% induction, only ∼5% of cells produced visible pili, rising steadily to WT levels at 0.3–0.4% arabinose. Similarly, the number of pili per cell in a 30-second window scaled with induction (Figure 2C): uninduced cells rarely produced pili (mostly 0, occasionally 1–2), while higher PilA levels restored distributions similar to WT. To better understand the mechanism that correlates pilin abundance to pilus number, we specifically analyzed the lengths of individual pili (Figure 2D). At 0% induction, observable pili were very short (<200 nm). The median lengths increased progressively with induction and reached WT values at 0.4% arabinose. These results establish that the abundance of PilA protein in the cell is a key determinant of pilus dynamics.

### Pilin abundance sets the maximum length of pilus fibers but not the count of pili

The sharp decrease in pilus length at low PilA levels prompted us to ask whether the apparent reduction in piliated cells under low induction was a true biological effect or an artifact of fluorescence microscopy resolution (where pili <200 nm would be undetectable). To resolve this at the molecular level, we employed computer simulations of pilus extension dynamics. Following our previous modeling framework [17, 48], we built a simulation that combines Brownian dynamics for the diffusion of PilA monomers in the inner membrane with a stochastic model of the PilB extension motor. PilB binds to the pilus base for random durations following an exponential distribution (median ∼2–4 seconds) [48], during which it can drive extension; monomer addition is then rate-limited by the ATPase’s cycle time (∼3 ms per PilA subunit under saturating conditions). In discrete 1 ms time steps, the model evaluates whether PilB is primed for a cycle, and whether a diffusing PilA monomer is close enough to the extension complex to be incorporated into the growing fiber. If both conditions are met, a monomer is added; otherwise, extension pauses for that step.

We then varied PilA membrane concentrations from 10% to 130% of WT levels and performed simulations for 10,000 individual pili per condition. Maximum and median pilus lengths increased with PilA availability, closely matching our experimental data (Figure 2E). At 10% PilA, the median length fell below 100 nm and medians exceeded 200 nm only above ∼30% PilA, the practical detection threshold in our microscopy setup. Based on this observation, we calculated the ratio of pili that were longer than 200 nm for each concentration of PilA (Figure 2F). As expected, the fraction of simulated pili longer than 200 nm rose from <10% at the lowest PilA concentration to ∼80% at WT levels.

These results explain our experimental pattern: at low PilA levels, the number of pili produced per cell does not decrease, the underlying count remains similar (governed by the availability of T4P complexes). However, a larger proportion of pili are too short to be visible by fluorescence microscopy. Thus, PilA abundance primarily controls pilus length (setting maximum and median values), while the number of pili per cell is largely independent of PilA levels. Consequently, a significant fraction of pili at low PilA levels remain too short to be observed, revealing a hidden subpopulation of filaments within the population. This also shows that controlling pilin abundance provides a reliable tool to tune median pilus length across a population without altering pilus count.

### Low pilin abundance forces extension into transient idle states that limit extension velocity

Motivated by our finding that PilA induction can reliably set the median pilus length, we sought to determine how pilin abundance modulates length at the molecular level. Pilus length is the product of extension velocity and the total duration of extension events. We therefore examined whether pilin levels influence one or both parameters.

Analysis of extension duration across induction conditions showed relatively little variation in the median time pili spent extending between 0.2% and 0.4% arabinose compared to WT levels (Figure 3A). Only at 0.1% induction did we observe a moderate reduction of 30–40%, with median extension times dropping to approximately 2 seconds. In contrast, extension velocity of individual pili increased steadily and significantly (by 77%) from 0.2% to 0.4% arabinose, with minimal change between 0.1% and 0.2% (Figure 3B). This velocity increase closely matches the 88% rise in median pilus length over the same range. These trends indicate that pilin abundance primarily affects extension velocity rather than the total duration of motor engagement (extension time). Our biophysical simulations recapitulated this pattern, showing a similar steady rise in extension velocity with increasing PilA levels that aligned well with the experimental measurements (Figure 3C). Importantly, retraction velocity remained constant across all conditions, confirming that monomer abundance specifically influences the extension mechanism and does not affect retraction (Figure 3D).

**Figure 3.**
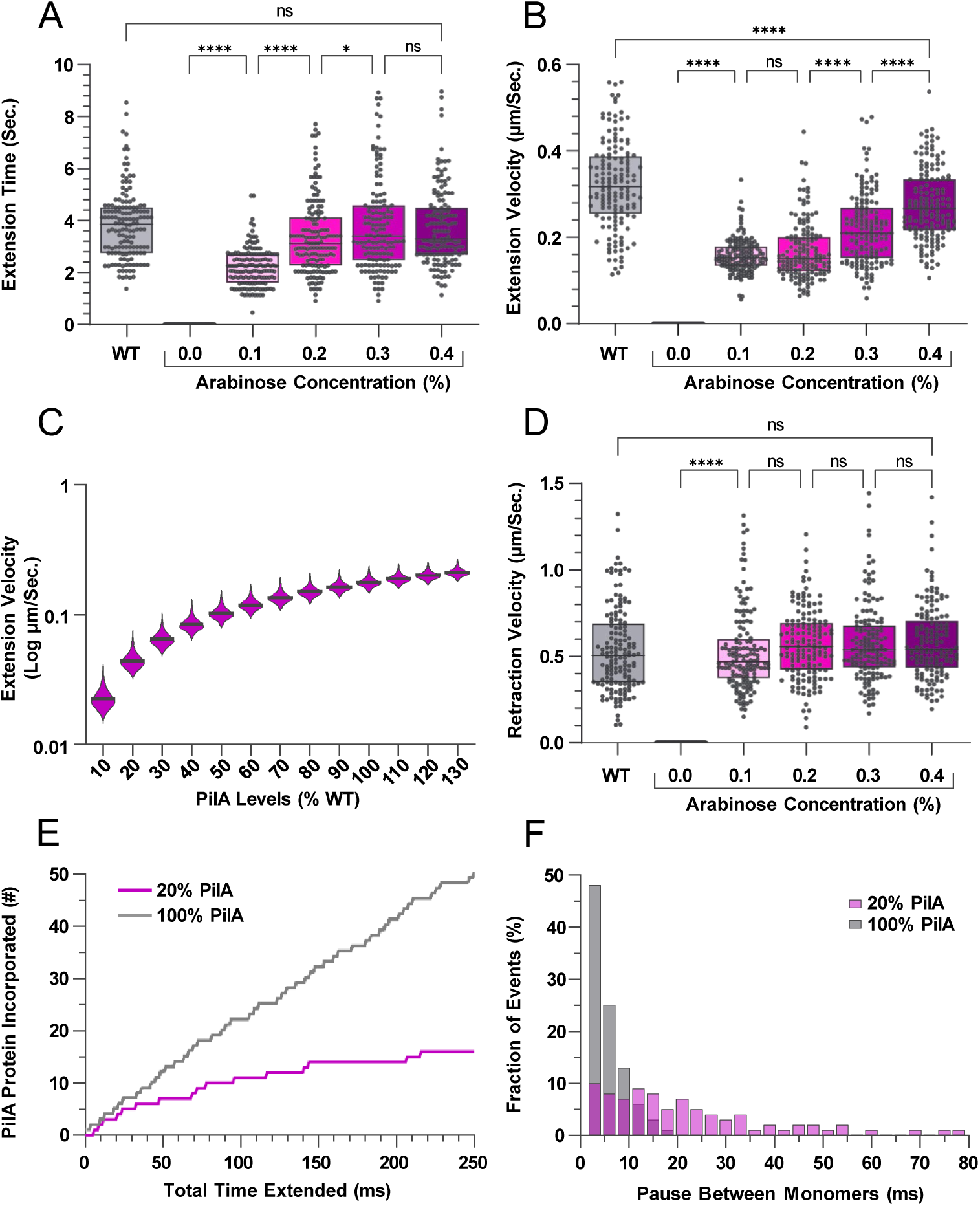
Changes in pilin levels modulate the extension velocity but not extension duration. **A)** Total duration individual pili spent extending until maximum length was reached as a function of PilA titration (*P_bad_*::*pilA* in a *pilA* background). **B)** Extension velocity of individual pili as a function PilA titration (same strain as in panel A). **C)** Extension velocity for 10,000 simulated pilus extension events as a function of PilA levels in the inner membrane. Violin plots show the median (horizontal line) and 5% to 95% percentiles. **D)** Retraction velocity of individual pili as a function of PilA titration (same strain as in panel A). **E)** Number of PilA proteins incorporated into the pilus fiber for 20% (purple) and 100% (gray) levels of total PilA from biophysical simulations. **F)** Percentage of events as a function of dwell times between individual PilA incorporations for 20% (purple) and 100% (gray) PilA levels. **A,B,D)** Boxes show median and 25th–75th percentiles of N = 150 pili from three independent biological replicates. Statistical tests were performed using one-way ANOVA: (ns) not significant, P > 0.05; (*) P < 0.05; (**) P < 0.01; (***) P < 0.001; (****) P < 0.0001.

To uncover the molecular basis for the velocity dependence on PilA, we examined the simulation trajectories in greater detail (Figure 3E). When focusing on a short 250 ms window of extension at 100% PilA levels, the time between consecutive monomer incorporations was dominated by the ATPase cycle time (∼3 ms, corresponding to ∼300 nm/s velocity in WT, assuming 1 nm per monomer), as expected. However, occasional longer intervals between incorporations were present. At 20% PilA, these delays became much more pronounced, with clear extended gaps between monomer additions. Histograms of the dwell times between individual incorporation events confirmed this shift (Figure 3F). At 100% PilA, roughly half of the intervals were limited by the ATPase cycle time, and no pauses exceeded 20 ms. At 20% PilA, only about 10% of events matched the ATPase cycle time, while pauses showed a broad distribution extending up to ∼80 ms. These results demonstrate that extension velocity is ultimately limited by the local availability of PilA monomers in the inner membrane. When pilin levels are low, monomers are not always present at the extension complex precisely when the ATPase is ready for the next cycle. This forces the extension machinery into transient idle states, reducing the effective incorporation rate and thereby constraining overall velocity.

### PilSR Regulates Pilus Dynamics Through Modulation of Pilin Levels

Our finding that pilin monomer abundance limits pilus extension velocity suggests that the PilSR two-component system regulates T4P dynamics by controlling PilA levels. To determine whether PilSR exerts any additional, direct effects on pilus dynamics independent of PilA abundance, we repeated the inducible PilA titration experiment in a *pilA pilSR* background and quantified the same key parameters: number of pili per cell (Figure 4A), the length of individual pili (Figure 4B), the extension duration (Figure 4C), and the extension velocity (Figure 4D). Across all four parameters, the trends with increasing arabinose induction were nearly identical to those observed in the *pilA* background alone. The number of pili per cell and median pilus lengths increased progressively with induction, reaching WT levels at ∼0.3–0.4% arabinose. Extension duration showed a similar plateau between 0.2% and 0.4% arabinose, with a comparable 30–40% reduction at 0.1% induction. Extension velocity also increased with induction, though the pattern differed slightly: unlike the *pilA* background (where velocity was flat between 0.1% and 0.2% before rising), velocity in the *pilA pilSR* background increased steadily from 0.1% to 0.4% arabinose.

**Figure 4.**
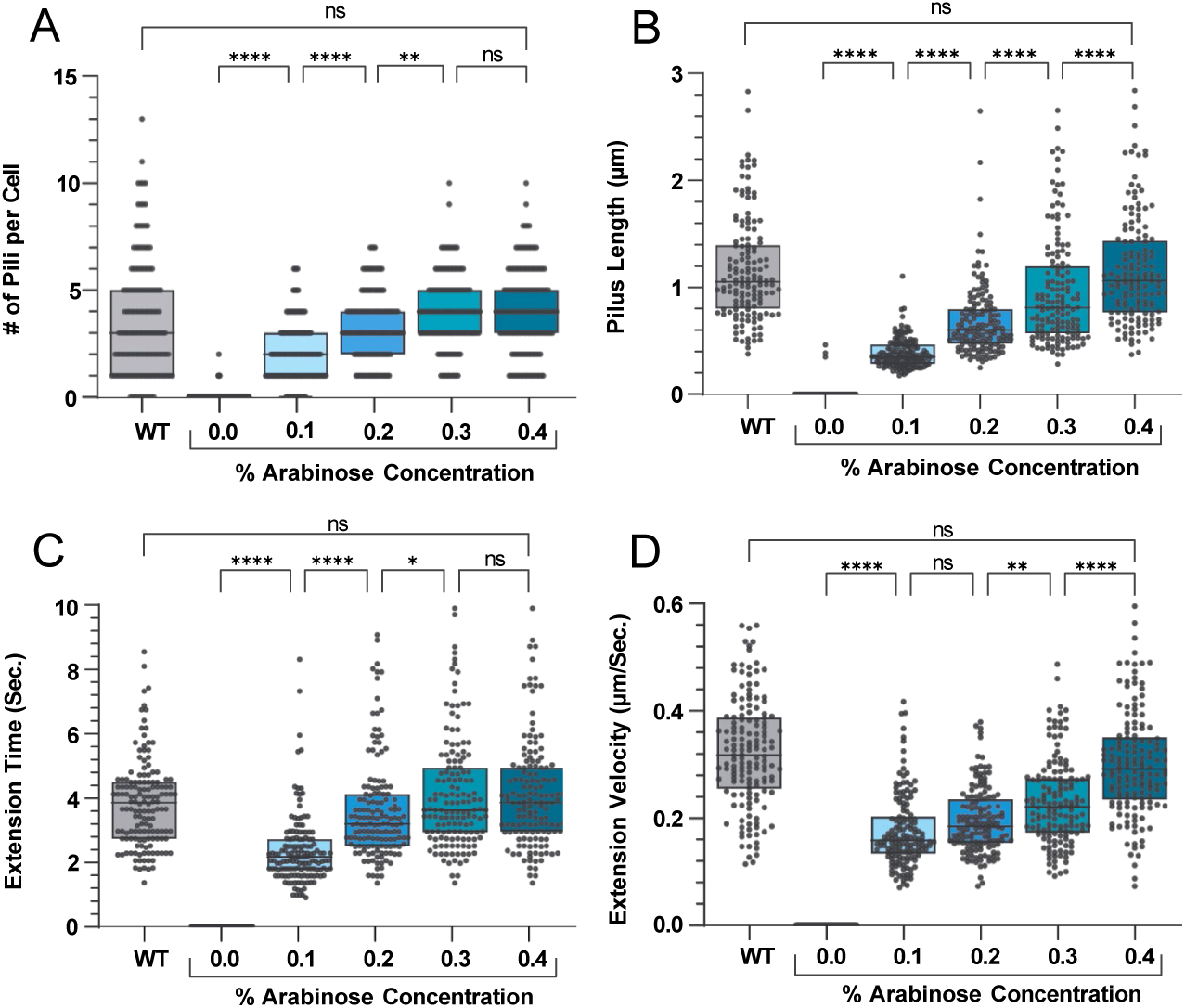
Pilus dynamics are not affected by PilSR directly. **A)** Total number of individual pili in a 30 s window in single cells as a function of PilA titration (*P_bad_*::*pilA* in a *pilApilSR* background). **B)** Maximum pilus extension lengths for individual pili as a function of PilA titration (same strain as in panel(A). **C)** Total duration individual pili spent extending until maximum length was reached as a function PilA titration (same strain as in panel (A). **D)** Extension velocity of individual pili as a function PilA titration (same strain as in panel (A). **A - D)** Boxes show median and 25th–75th percentiles of N = 150 cells (A) or pili (B-D) from three independent biological replicates. Statistical tests were performed using one-way ANOVA: (ns) not significant, P > 0.05; (*) P < 0.05; (**) P < 0.01; (***) P < 0.001; (****) P < 0.0001.

These highly similar behaviors indicate that deleting PilSR does not substantially alter the relationship between PilA abundance and pilus dynamics. Collectively, the data demonstrate that PilSR modulates T4P dynamics primarily through control of pilin levels, with no strong evidence for additional direct mechanisms acting on extension or retraction.

### Pilus length limits TFP-mediated virulence behaviors

Our observation that a substantial fraction of pili are too short to be detected by fluorescence microscopy suggests that they might also be too short for functional contribution, for example to different virulence traits. This prompted us to investigate how pilus length influences T4P-dependent virulence traits in *P. aeruginosa*. To test this, we used the inducible PilA system to titrate pilus length and assessed twitching motility, surface sensing, phage infection, and biofilm formation across a range of arabinose concentrations.

#### Twitching Motility

Twitching motility requires pili to make physical contact with a surface and retract to pull the cell forward. In a standard stab-agar twitch plate assay, twitching zone diameter increased gradually with arabinose induction, mirroring the trends in pilus dynamics (Figure 5A). Wild-type twitching levels were reached only at 0.5% arabinose, higher than the 0.3–0.4% needed for wild-type PilA protein levels. This delay is consistent with pili at 0.4% being approximately 15% shorter than in WT cells. Further increasing arabinose to 0.6% produced no additional gain in twitching. In the *pilSR* background, twitching increased more rapidly with induction and plateaued at 0.3% arabinose, with maximum twitching being 20% higher than in the WT background (Figure 5B). These results demonstrate that pilus length directly limits twitching capacity: shorter pili reduce the efficiency of surface contact and retraction. The enhanced twitching in the *pilSR* strain further suggests that PilSR negatively regulates twitching motility through targets outside the T4P system, independent of its effect on PilA levels.

**Figure 5.**
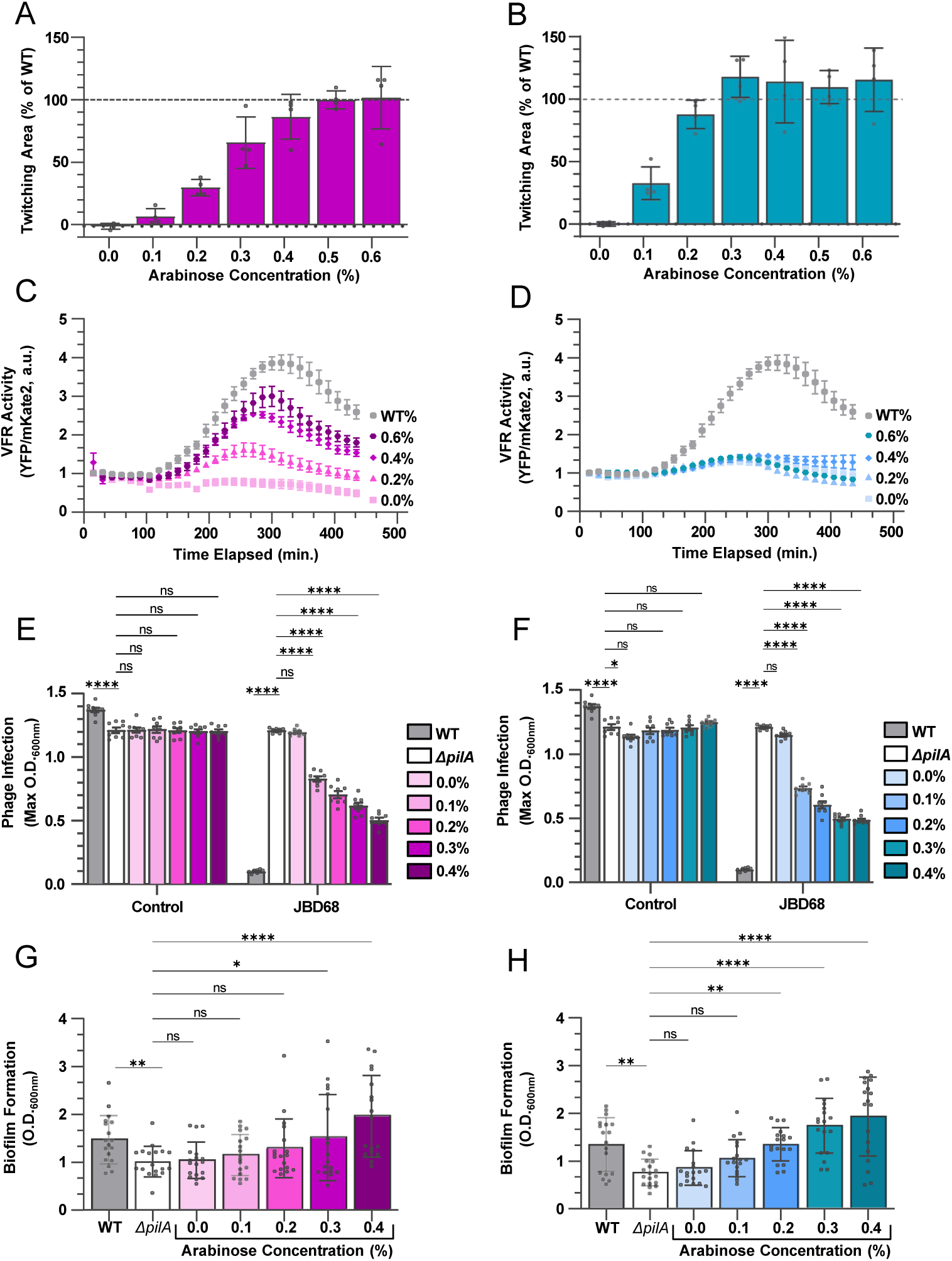
The length of pili governs T4P-dependent virulence traits. **A,B)** The area of twitching motility as percentage of WT on 1% LB agar plates as a function of PilA titration (*P_bad_*::*pilA*) in *pilA* (A) and *pilA pilSR* (B) backgrounds. The horizontal dotted line indicates the mean twitching area of the WT strain under identical conditions. Bars represent mean ± SD of three biological replicates (indicated as black dots). **C,D)** Surface sensing activity over time for cells on a 1.5% agarose pad. Surface sensing was measured by the activity of the transcription factor Vfr using the fluorescent reporter PaQa as a function of PilA titration (*P_bad_*::*pilA*) in *pilA* (C) and *pilA pilSR* (D) backgrounds. Data points represent mean ± SEM of three biological replicates. **E,F)** Maximum bacterial growth (O.D. = optical density) after infection with the pilus tip-binding phage JBD68 as a function of PilA titration (*P_bad_*::*pilA*) in *pilA* (E) and *pilA pilSR* (F) backgrounds (see Supplementary Information for individual growth curves). Bars represent mean OD_600_ ± SEM (n = 8 technical replicates across 4 biological replicates). **G,H)** Biomass of biofilms grown on liquid-submerged pegs as a function of PilA titration (*P_bad_*::*pilA*) in *pilA* (G) and *pilA pilSR* (H) backgrounds. Bars represent mean OD_600_ ± SD of N = 18 technical replicates across three biological replicates. **G-H)** Statistical tests were performed using unpaired t-test. **E – F)** Statistical tests were performed using one-way ANOVA: (ns) not significant, P > 0.05; (*) P < 0.05; (**) P < 0.01; (***) P < 0.001; (****) P < 0.0001.

#### Surface Sensing

Similar to twitching, surface sensing in *P. aeruginosa* relies on pili detecting mechanical cues from substrate contact, triggering cAMP production and Vfr-dependent gene expression via the Pil-Chp pathway [19, 20, 33, 36, 62, 63]. We monitored this response using the PaQa transcriptional fluorescent reporter for Vfr activity on 1.5% agarose pads over a 7-hour time course. WT cells showed a progressive increase in Vfr activity, peaking at approximately four-fold upregulation relative to liquid-grown cells after five hours (Figure 5C). At 0% arabinose, the response was completely abolished. Increasing induction gradually restored the response, with amplitudes rising from 1.5-fold at 0.2% to 2.5-fold at 0.4% arabinose. Similar to twitching, WT PilA protein levels (0.3–0.4%) did not fully restore WT sensing; even at 0.6% induction, the response reached only 3-fold. In the *pilSR* background, surface sensing was abolished across all inducer concentrations (Figure 5D). These findings indicate that both pilus length and the ability to modulate PilA levels via PilSR are essential for efficient surface sensing through the Pil-Chp pathway.

#### Phage Infection

Unlike twitching and surface sensing, phage infection does not require pili to contact a solid substrate. Instead, many phages bind along the pilus fiber and use retraction to reach the surface of the cell. We tested infection efficiency using phage JBD68 (which targets the tip protein FimU) in a liquid growth assay, monitoring optical density (OD) over 12 hours after infection (Supplementary Figure 2). Infected WT cultures showed an initial rise in OD followed by a sharp decline due to lysis, with maximum OD dropping to ∼0.1 compared to uninfected controls (∼1.4) or *pilA* (∼1.2) (Figure 5E, Supplementary Figure 2A,B). Increasing PilA induction caused a progressive decrease in maximum OD, indicating more efficient phage infection with longer pili. The pattern was similar in the *pilSR* background, with no significant difference across conditions (Figure 5F, Supplementary Figure 2C,D). These results show that pilus length enhances phage infection efficiency, likely by increasing the probability of productive tip binding and retraction, but PilSR does not appear to play a major role in this process.

#### Biofilm Production

Lastly, T4P contribute to biofilm formation through adhesion and microcolony spreading, however, the mechanisms of how T4P dynamics contribute to more robust biofilms remains largely unknown [11, 32, 62, 64]. We quantified the formation of biofilm biomass on peg lids in a 96-well microtiter format after 18 hours of growth, using crystal violet staining and acetic acid solubilization of bound dye (Figure 5G). The *pilA* control produced ∼40% less biofilm mass than WT and gradual PilA induction restored biofilm biomass to WT levels by 0.4% arabinose (Figure 5G). The *pilSR* background showed a similar restoration pattern (Figure 5H). These data indicate that robust biofilm formation depends on pilus length, with shorter pili limiting overall biomass accumulation.

## Discussion

Type IV pili (T4P) enable bacteria to interact with their environments through dynamic cycles of extension and retraction. Much of the literature has focused on retraction as the critical rate-limiting step in T4P function, highlighting the high forces (>100 pN per pilus) generated by retraction ATPases (PilT/PilU) that power twitching motility, surface sensing, phage uptake, natural transformation, and biofilm maturation [9, 13, 17–19, 29–36, 51–57]. Extension on the other hand is typically seen as a mere preparatory step. Here, we challenge this perspective by showing that extension dynamics, specifically pilus length governed by PilA monomer availability, represent a critical overlooked bottleneck that constrains all retraction-enabled virulence traits.

We demonstrated that PilA levels vary ∼10-fold between cells in natural populations despite PilSR regulation. Our inducible titration system, combined with biophysical simulations, revealed that PilA abundance primarily tunes extension velocity and pilus length, not pilus count per cell. At low PilA concentrations, diffusion-limited monomer supply in the 2D inner membrane increases waiting times for PilA to reach the PilB extension complex, forcing transient idle states where PilB is primed but cannot incorporate subunits. This slows net extension velocity and caps length, while retraction velocity remains unaffected. Because extension events terminate stochastically via PilB unbinding, the resulting pilus lengths follow an exponential distribution inherently biased toward short filaments [13, 48]. This bias produces a large hidden subpopulation of non-contributing pili that cannot meaningfully interact with substrates, host cells, or phages. Binding to DNA, phages, or surfaces during twitching requires pili to extend sufficiently far, often beyond the cell’s extracellular polysaccharide coat or to reach a distant target, much like a fishing line must be long enough to reach the water to catch fish.

For example, the mean pilus length at low PilA induction (20%) is ∼160 nm and increases 4.7-fold to ∼750 nm for WT-like cells (100%, simulations). Assuming the functional threshold needed for effective substrate interaction is similar to the mean length of T4P in WT cells (750 nm), this increase in mean length by 4.7-fold raises the fraction of contributing pili (the long tail of the exponential distribution) from ∼0.9% to 37%, yielding approximately 40-fold more pili capable of contributing to adhesion or motility (Supplementary Figure 3). This represents a strong non-linear lever that boosts the ratio of contributing pili disproportionately to the increase in pilin abundance. Achieving the same gain by increasing pilus count alone (length distribution unchanged at 160 nm) would require scaling total pilus number ∼40-fold, which is biologically challenging. Cells typically maintain only ∼5 pilus machines per cell (each comprising ∼50 subunits of PilCMNOPQ, plus ∼10× more PilB/T/U hexamers), limited by inner-membrane space and assembly energetics. Scaling to 40-fold more machines would demand enormous protein synthesis, membrane insertion, and coordinated assembly, far exceeding the modest cost of elevating PilA from ∼2000 molecules (20% induction) to ∼10,000 (WT levels). Moreover, such expansion would likely overwhelm downstream signaling control via Pil-Chp, PilZ, and other regulators, risking dysregulation and interference with other membrane functions [19, 20, 33, 36–38, 62, 63, 65, 66]. In contrast, increasing PilA abundance is transcriptionally simple, energetically modest, and avoids these bottlenecks, enabling precise, low-cost shifts in the contributing fraction via the exponential tail of the distribution. This makes length regulation far more efficient, and likely biologically favored, than count regulation for enhancing T4P-mediated virulence traits, and presents another strong non-linear functional lever in the efficiency of increasing the ratio of contributing pili relative to an increase in pilus machine count.

Our experiments support this conclusion across T4P-mediated behaviors: longer pili consistently restored function more effectively than equivalent changes in pilus count alone would imply. Twitching motility and surface sensing (via Pil-Chp/Vfr) showed the strongest length dependence, with PilSR providing additional independent regulation, negative for twitching (likely via unknown targets such as surfactants or adhesion components) and essential for surface sensing. Phage infection and biofilm formation scaled primarily with length and showed little PilSR sensitivity, suggesting partial decoupling from mechanosensing pathways.

We like to note a recent study in *Vibrio cholerae* that reported lowering of PilA levels increased pilus number, in contrast to our observations in *P. aeruginosa* [67]. These complementary findings highlight how PilA titration effects can be context-dependent (species, mutant background) and reinforce that monomer availability critically shapes T4P dynamics and functional output. It will be interesting to uncover how context-dependent and other confounding biological factors regulate T4P dynamics and their associated virulence traits in future studies.

Together, our study reframes the importance and functional consequences of T4P dynamics: extension velocity, limited by diffusion-driven monomer supply and idle states, governs the effective fraction of pili that contribute to function and thus constrains retraction-powered virulence. The exponential length distribution, combined with ∼10-fold cell-to-cell variation in PilA levels, generates substantial phenotypic heterogeneity in pilus length within a clonal population. This heterogeneity enables bet-hedging: subpopulations with predominantly short pili may persist under conditions where long pili are costly or risky (e.g., phage exposure or energy limitation), while those with longer pili are poised for rapid colonization when opportunities arise. Such diversity increases the overall resilience of the population to fluctuating environments without requiring every cell to commit to a single strategy. This mechanism may extend to other T4P pathogens, where length heterogeneity influences pathogenesis. Future studies on length thresholds in vivo and regulatory feedback loops will further clarify how extension emerges as a governing step in T4P biology.

## Acknowledgements

We would like to thank Joanne Engel and Yuki Inclan for the gift of the polyclonal PilA antibody, Karen Maxwell and Veronique Taylor for the gift of phage JBD68, Joseph Sorg and Morgan Osborne for training and use of the LiCOR Odyssey M. We would like to thank Joseph Sanfilippo and the entire Koch lab for stimulating discussion, and the Department of Biology at Texas A&M for its supportive environment.

This work was supported by grant R35GM155280 from the National Institute of Health and startup funds from the College of Arts and Sciences, Division of Research, and Department of Biology at Texas A&M University to M.D.K.

## Author contributions

Z.M., J.R., and M.D.K. designed research. Z.M. and M.D.K. wrote the manuscript. Z.M., J.R., A.O.Y, and M.D.K. provided reagents and constructs. Z.M., T.P., N.C, and S.D. performed experiments and analyzed data.

## Methods and Protocols

### Strains and growth conditions

*P. aeruginosa* PAO1 was grown in liquid lysogeny broth (LB) Miller (Difco) in a floor shaker, or on LB Miller agar (1.5% Bacto Agar), on Vogel-Bonner minimal medium agar (200 mg/L MgSO4 7H2O, 2 g/L citric acid, 10 g/L K2HPO4, 3.5 g/L NaNH4HPO4 4 H2O, and 1.5% agar), and on no-salt LB agar (10 g/L tryptone, 5 g/L yeast extract, and 1.5% agar) at 30 °C (for cloning) or at 37 °C. *E. coli* S17 was grown in liquid LB Miller (Difco) in a floor shaker and on LB Miller agar (1.5% Bacto Agar) at 30 °C or at 37 °C. Arabinose-supplemented media were prepared by adding filter-sterilized 4% (w/v) arabinose stock solution to LB or autoclaved medium to achieve the desired final concentration. Antibiotics were used at the following concentrations: 100μg/mL or 200 μg/mL carbenicillin in liquid (300 μg/mL on plates) or 10 μg/mL gentamycin in liquid (30 μg/mL on plates) for *P. aeruginosa*, and 100 μg/mL carbenicillin in liquid (100 μg/mL on plates) or 30 μg/mL gentamycin in liquid (30 μg/mL on plates) for *E. coli*. Western blotting medias were formulated as follows: 4x loading Dye (4ml of Glycerol, 2.5 mL of 1M Tris-HCL pH6.8, 20 mg Bromophenol blue, 2 mL β-mercaptoethanol, 0.8 g SDS, 1.5 ml H20), 10x Tris-Buffer Saline ph7.6 (24.23g of Tris, 80g NaCl, adjusted to 1 l wtih H2O), 1x Tris-Buffer Saline with Tween-20 (1 ml Tween 20, 100 ml of 10x TBS, adjusted to 1 l withH2O), 1x Running Buffer (12 g Tris, 57.6 g of Glycine, 4g SDS, adjusted to 4 l with H2O), Blocking Buffer (2.5g of powdered milk, 50 ml of 1x TBS).

### Generation of arabinose-inducible pilA construct (pMK119)

The mini-Tn7 delivery vectors pMK119 harboring either a native copy of pilA or the A86C allele were constructed by Gibson assembly. The backbone was obtained by linearizing pUC18-mini-Tn7T-LAC with Eco53kI and NsiI restriction enzymes. The fragment containing *araC* and *P_bad_* was PCR amplified from the iRFP670 plasmid [68] using primers pMK119_f1.For/Rev and digested with AvrII. The *pilA* fragments were PCR amplified using primers pMK119_F2.For/Rev from genomic DNA. All fragments were assembled using NEBuilder HiFi DNA Assembly Master Mix (New England Biolabs) according to the manufacturer’s protocol. Assembly products were electroporated into electrocompetent *E. coli* S17: 50 µL competent cells were mixed with the assembly product, transferred to a chilled electroporation cuvette, and pulsed. Immediately after electroporation, 1 mL LB was added, and cells recovered at 37°C with shaking for 1–1.5 h. Cells were concentrated by centrifugation, resuspended in 100 µL LB, and plated on LB agar supplemented with 100 µg/mL carbenicillin. Plates were incubated overnight at 37°C. Constructs were verified by colony PCR and confirmed by Sanger sequencing using primers pTn7_Ver1/2. Verified strains were grown overnight in LB + 100 μg/ml carbenicillin, and glycerol stocks (25% v/v) were prepared and stored at −80°C.

### Chromosomal integration of pMK119

Plasmids pMK119 with either the native of A86C allele of *pilA* and helper plasmid pTNS2 were purified using the Qiagen miniprep kit. Electrocompetent recipient strain were prepared by growing cultures overnight in LB, diluting 1:1000 into fresh LB, and harvesting at early stationary phase. Cells were washed three times in ice-cold 300 mM sucrose and resuspended in 50 μl of the same buffer. For each electroporation, 50 μl competent cells were mixed with 300 ng each of pMK119 and pTNS2 plasmid DNA, electroporated, and recovered in 1 ml LB at 37°C with shaking for 1–1.5 h. Cells were concentrated, plated on LB agar + 10 μg/ml gentamycin, and incubated overnight at 37°C. Successful insertion at the attTn7 site was confirmed by colony PCR using primers pTN7_Ver1/2, which produce a diagnostic band shift. Confirmed integrands were grown overnight in LB + 10 μg/ml gentamycin and stored as 50% glycerol stocks at −80°C.

### Generation of *pilSR* recipient strain

In-frame deletions were generated using allelic exchange with the suicide vector pEXG2 following [69]. Flanking regions upstream of pilS and downstream of piLR were amplified with PCR using primers pilS_P1/P2 and pilR_P3/P4. Fragments were joined by overlap extension PCR using the outermost primers, digested with HindIII-HF, and ligated into HindIII-digested pEXG2. Constructs were transformed into *E. coli* S17, verified by PCR and Sanger sequencing with pEXG2_Ver1/Ver2 primers, and introduced into PAO1 by conjugation. For mating, 1.5 ml donor containing the vector were grown to OD 0.5. The PA01 parental strain was grown overnight, and 0.5 ml culture was diluted 1:2 into fresh LB and incubated for 3 hours at 42 °C. Both cultures were concentrated into 100 μl and spotted onto an LB agar plate and incubated overnight at 30 °C. The puddle was scrapped off, resuspended into 150 μl PBS and spread on VBMM + 30 μg/ml gentamicin at 37°C for 24 h. For counter-selection, single colonies from the VBMM plate were struck onto NSLB and incubated for 24 hours at 30 °C. Colonies from the NSLB plate were screened for the correct deletion mutation using PCR amplification with the flaking primers and confirmed using sanger sequencing.

### Construction of *PpilA*::*yfp* transcriptional reporter plasmid

The *P_pilA_*::*yfp* reporter plasmid was constructed using the established *P_PaQa_*::*yfp P_rpoD_*::*mKate*2 dual-reporter vector that includes the constitutive *P_rpoD_* promoter for normalization [19]. The vector was linearized by digestion with XhoI and BsiWI. The native *pilA* promoter region and *yfp* coding sequence were PCR-amplified as separate fragments using primers PpilA_F1.For/REV (*yfp*) and PpilA_F2.For/REV (*pilA* promoter). All fragments were assembled using NEBuilder HiFi DNA Assembly Master Mix (New England Biolabs) according to the manufacturer’s protocol. electroporated into electrocompetent *E. coli* S17, and 50 µL competent cells were mixed with the assembly product, transferred to a chilled electroporation cuvette, recovered in 1 mL LB and grown at 37°C with shaking for 1–1.5 h. Cells were then concentrated by centrifugation, resuspended in 100 µL LB, and plated on LB agar supplemented with 100 µg/mL carbenicillin. Plates were incubated overnight at 37°C. Successful assembly was confirmed by colony PCR using primers PpilA_Ver1/2, which flank the insertion site and produce a diagnostic band shift. Verified clones were grown overnight in LB + 100 μg/ml carbenicillin at 37°C with shaking, mixed 1:1 with 50% (v/v) glycerol, and stored as frozen stocks at −80°C.

This reporter plasmid was then isolated using a Qiagen miniprep kit and introduced into recipient strains using electroporation. Electrocompetent cells were prepared by growing cultures overnight in LB, diluting 1:1000 into fresh LB, harvesting at early stationary phase, washing three times in ice-cold 300 mM sucrose, and resuspending in 50 μl of the same buffer. For each electroporation, 50 µL competent cells were mixed and with 100ng of donor plasmid, transferred to a chilled cuvette, electroporated, recovered in 1 mL LB at 37°C shaking for 1–1.5 h. Cells were concentrated, resuspended in 100 µL LB, and plated on LB agar containing 300 µg/mL carbenicillin. Plates were incubated overnight at 37°C. Successful plasmid maintenance was confirmed by colony PCR using priemrs PpilA_Ver1/2. Confirmed transformants were grown overnight in LB + 200 μg/ml carbenicillin and stored as 25% glycerol stocks at −80°C.

### Twitch Plate Assay

LB broth supplemented with 1% (w/v) agar was autoclaved and allowed to cool to approximately 50–60°C. Specified concentrations of L-arabinose were added to separate 100 mL aliquots of the cooled molten agar, mixed thoroughly, and poured evenly into Petri dishes. Plates were allowed to solidify and were stored overnight at room temperature to dry. Bacterial strains were recovered from frozen glycerol stocks by streaking onto standard LB agar plates (1.5% agar) and incubating overnight at 37°C. The following day, single colonies from these plates were used to inoculate designated areas on the surface of the arabinose-supplemented 1% LB agar plates by spotting through the agar layer to ensure contact with the agar-plastic interface. Plates were incubated at 37°C for 3 days to permit development of twitching motility zones at the agar-Petri dish interface. Following incubation, the agar was carefully removed and discarded. A 1% (w/v) crystal violet solution was added to stain the biofilm matrix on the Petri dish surface for 10–15 min. Excess stain was gently rinsed off with deionized water, and plates were air-dried. The total area of twitching motility zones was measured using digital imaging. Mutant twitching zones T_M_ were normalized to WT (T_WT_, positive control) and *pilA* (T_pilA_ negative control) to yield percentage of WT twitching according to (T_M_ – T_pilA_) / (T_WT_ – T_pilA_).

### Phage Infection Growth Assay

Phage infection assays were carried out in 96-well plates liquid culture formats as described previously [70–72]. In brief, Bacterial strains were inoculated into LB broth and grown overnight at 37°C with shaking (250 rpm). The following day, overnight cultures were diluted 1:333 into 2 mL of fresh LB medium supplemented with the appropriate concentration of L-arabinose and grown at 37°C with shaking until an optical density OD₆₀₀ = 0.5 was reached. These mid-log phase cultures were then diluted 1:100 into 2 mL of fresh arabinose-supplemented LB medium for a concentration of 5*10^6^ CFU/mL. For the infection assay, 25 µL of each phage dilution (total PFU: 3.87*10^6^ PFU/mL) was added to designated wells of a clear flat-bottom 96-well microtiter plate. Subsequently, 225uL (Total CFU: 1.12*10^6^ CFU/mL) of the diluted bacterial cultures were added to designated wells, resulting in a final volume of 250 uL per well (MOI ∼3.44). Bacterial growth was monitored by measuring OD600 every 15 min overnight at 37C with continuous orbital shaking in a microplate reader.

### Biofilm Assay

Biofilm assays were carried out in 96-well plates liquid culture formats as described previously [73]. In brief, Bacterial strains were inoculated into LB broth and grown overnight at 37°C with shaking (250 rpm). The following day, overnight cultures were diluted 1:333 into 2 mL of fresh LB medium supplemented with the appropriate concentration of L-arabinose and grown at 37°C with shaking until an optical density OD₆₀₀ = 0.5 was reached. These mid-log phase cultures were then diluted 1:100 into 2 mL of fresh arabinose-supplemented LB medium. 180 µL of the resulting dilution were dispensed into designated wells of a 96-well microtiter plate. A Nunc-Immuno™ TSP 96-peg lid was placed onto the plate, and biofilms were allowed to form during overnight incubation at 37 °C. The following day, the peg lid was gently washed by immersion in a 96-well plate containing 180 µL of sterile phosphate-buffered saline (PBS) per well. Biofilms were then stained by transferring the peg lid to a separate 96-well plate containing 180 µL of crystal violet solution per well and incubating for 15 min at room temperature. Excess stain was removed by washing the peg lid in new 96-well plate containing 180 µL of PBS per well. For biofilm solubilization, the peg lid was transferred to a new 96-well plate containing 180 µL of 100% acetone per well and incubated for 15 min at room temperature. The peg lid was subsequently removed, and biofilm biomass was quantified by measuring the absorbance at 600nm using a microplate reader.

### Western Blots

Bacterial strains were inoculated into LB broth and grown overnight at 37°C with shaking (250 rpm). 800 µL of overnight culture was pelleted by centrifugation at 13,000 rpm for 2 min, the supernatant was discarded, and the pellet was resuspended in 800 µL of fresh LB medium. This washed culture was used to inoculate 4 mL of fresh arabinose-supplemented LB medium at a 1:333 dilution and grown at 37 °C with shaking until an optical density OD₆₀₀ = 0.8 was reached. Cells were harvested from 800–2400 µL of culture (adjusted based on protein expression levels) by centrifugation. Supernatants were removed, and pellets were washed by resuspension in 200 µL of phosphate-buffered saline (PBS) followed by centrifugation. Final pellets were resuspended in 20 µL of SDS sample loading buffer and heated at 95 °C for 20 min.

Protein samples (5 µL) and LI-COR Chameleon Duo protein ladder (0.8 µL) were loaded into 20% Mini-PROTEAN TGX precast gels (Bio-Rad). Electrophoresis was performed using the Mini-PROTEAN system (Bio-Rad) in pre-chilled running buffer with the gel apparatus surrounded by ice until the dye front reached the bottom of the gel.

Nitrocellulose membranes and filter papers were pre-equilibrated in transfer buffer for at least 1h prior to transfer. Proteins were transferred to nitrocellulose membranes using the Trans-Blot Turbo system (Bio-Rad) according to the manufacturer’s instructions. Following transfer, total protein was visualized using Revert™ 520 Total Protein Stain (LI-COR) for loading normalization. Membranes were washed in Tris-buffered saline (TBS), dried at 37 °C for 20 min, rehydrated in TBS, and rinsed in deionized water. Membranes were incubated in Revert 520 stain until protein bands were visible, then washed twice in Revert Wash Solution followed by a wash in deionized water. Total protein signal was imaged using an Odyssey CLX imager (LI-COR) in the 520 nm channel. Membranes were subsequently rinsed in ultrapure water for 5 min, destained using Revert Destaining Solution until background signal was cleared, and rinsed again in ultrapure water for 5 min while shaking.

For immunodetection, membranes were washed in TBS and blocked in blocking buffer for 1 h at room temperature with shaking. Membranes were then incubated overnight at 4 °C with shaking in primary antibody (rabbit anti-PilA polyclonal antibody, 1:50,000 dilution in blocking buffer). The following day, membranes were washed three times in TBST and incubated for 1 h at room temperature with shaking in IRDye-conjugated secondary antibody (1:50,000 dilution in blocking buffer). Membranes were washed three additional times in TBST and imaged using an Odyssey CLX imager (LI-COR) in the appropriate fluorescence channels. Band intensities for PilA were quantified and normalized to total protein signal obtained from Revert 520 staining.

### Fluorescence microscopy imaging

Fluorescence imaging was performed using a Nikon Ti2-Eclipse inverted microscope controlled by NIS-Elements software (Nikon), equipped with a 40x/0.95NA air and a 100x/1.45NA oil objective (Nikon), an ORCA-Fusion BT sCMOS camera (Hamamatsu), 488/514/561 nm lasers and appropriate filters and dichroic optimized for imaging of Alexa488, YFP, and mKate2 channels.

### Imaging of PaQa and PilA transcriptional reporters

Surface sensing assays were carried out in 96-well plates liquid culture formats as described previously [17, 19]. Bacterial strains were inoculated into LB broth with Carb100 and grown overnight at 37°C with shaking (250 rpm). The following day, cultures were diluted 1:333 into fresh LB medium supplemented with arabinose and grown to OD₆₀₀ = 0.5. 300 µL aliquots of cultures were transferred to microcentrifuge tubes, pelleted by centrifugation at 13,000 rpm for 2 min, and the supernatant was removed. Cell pellets were resuspended in 300 µL of fresh LB medium. 1 uL of each of the resuspended cultures were spotted onto the center of an agarose pad (1%) supplemented with the designated concentration of arabinose. Agarose pads were then inverted onto glass-bottom dishes for imaging. For PaQa, time-lapse images were acquired every 15min for a total duration of 7h. Each experiment consisted of four independent fields of view (FOV) of each agarose pad. FOVs containing approximately 10-40 cells were selected for imaging. Individual cells were segmented using a custom-written, threshold-based image analysis algorithm based on fluorescence from a constitutively expressed reporter (*P_ropD_*::*mKate*2) [17]. This segmentation approach enabled estimation of mean background and fluorescence intensities for both reporter channels. Transcriptional reporter activity was quantified as the ratio of YFP to mKate2 fluorescence. For each time point, the median fluorescence ratio for all cells was calculated. For the PilA transcriptional reporter, the imaging setup and analysis was identical but only one still image was taken.

### Imaging of Pilus Dynamics

Pilus dynamics microscopy assays were carried out agarose gel pad from liquid culture as described previously [48]. Bacterial strains were inoculated into L) broth and grown overnight at 37°C with shaking (250 rpm). The following day, overnight cultures were diluted 1:333 into fresh EZ Rich Defined Medium supplemented with the specified concentration of L-arabinose. Cultures grew at 37°C with shaking until mid-log phase, approximately OD_600_ = 0.5. 200 µL of each culture was transferred to a 1.5 mL microcentrifuge tube and 1uL of Alexaflour-488 maleimide dye (dissolved in anhydrous DMSO at 2.5ug/uL) was added. The mixture was incubated for 45 min at 37C in the dark. Labeled cells were gently washed twice to remove unbound dye by pelleting and resuspension in 200 µL of fresh EZ rich media. After the second wash, cells were finally resuspended in 50 µL of fresh EZ rich media. A 2 µL of the resuspended cells was spotted onto the center of an agarose pad, the pad was then inverted onto a glass-bottom dish. Time-lapse imaging of type IV pilus dynamics was performed every 200ms for 30s at minimal laser power to reduce phototoxicity and photobleaching. Single-cell TFP dynamics (extension, retraction, count, and length) were analyzed manually in Fiji/ImageJ.

