## Supplementary Information for "Type IV pilus length determines virulence by regulating a hidden subpopulation of non-contributing filaments"

### Supplementary Results

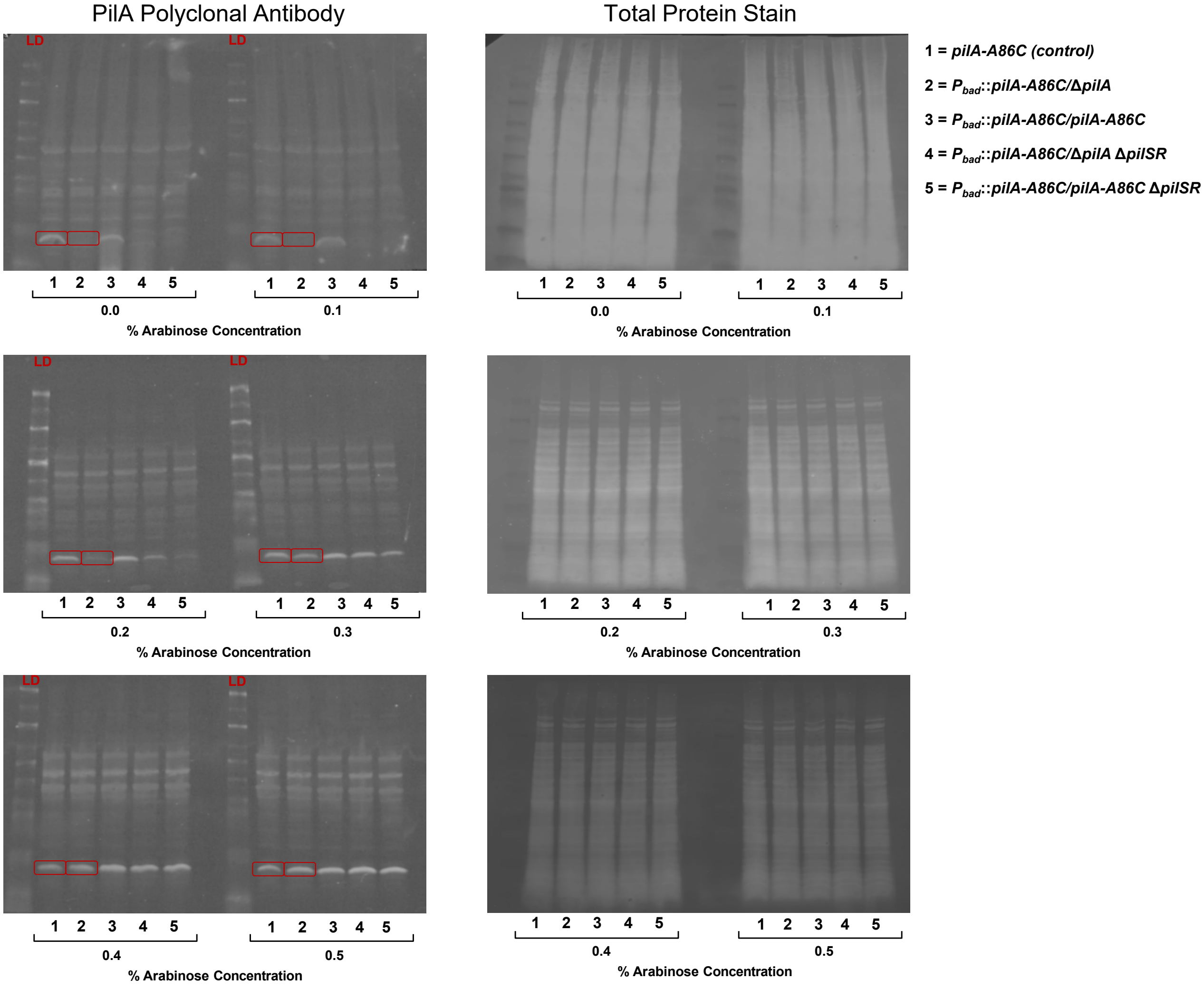

**Supplementary Figure 1. Representative western blot images for PilA protein levels showing levels of PilA protein targeted by PilA-antibody (left column) and corresponding total protein control (rigth column) with PilA titration (0 – 0.5% arabinose).** Molecular weight ladders (LD) are shown in the left most column for each treatment group (two groups per blot). Control WT (**1**) and *$P_{bad}$::pilA-A86C/ΔpilA* (**2**) are outlined with red boxes (Molecular weight ~15kDa) and were used for quantification in Figure 2A.

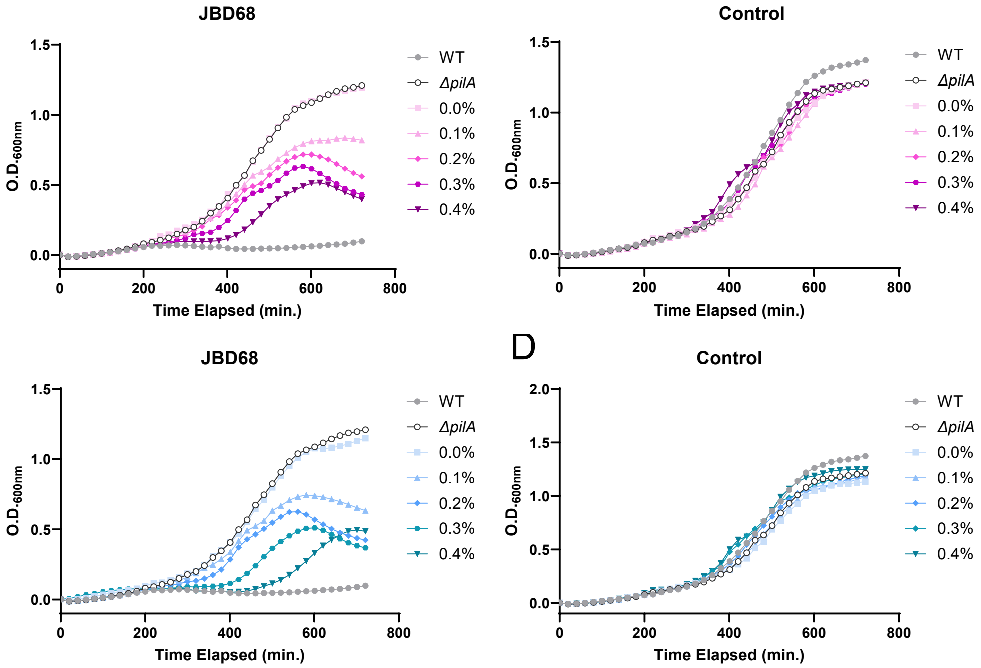

**Supplementary Figure 2. Bacterial growth curves after infection with the pilus tip-binding phage JBD68 as a function of PilA titration ($P_{bad}$::*pilA*) in Δ*pilA* (A,B) and Δ*pilSR* (C,D) backgrounds over a 12 hour time course.** Data points represent the mean of four biological replicates and two technical replicates each.

### Supplementary Methods

**Biophysical simulation of pilus extension and PilA monomer diffusion:** To investigate how PilA abundance in the inner membrane constrains pilus extension velocity and final length, we developed a Brownian dynamics model of monomer diffusion coupled to a stochastic extension motor, similar to [1, 2]. The inner membrane is modeled as a spherical surface of radius 0.3 µm (reflecting the geometry of the cell pole). PilA monomers (N particles) diffuse freely on this surface with a diffusion coefficient D = 0.22 $\mu m^2/s$ [2]. The extension motor PilB is positioned at a fixed reference point on the membrane (corresponding to the center of the cell pole). Monomers are incorporated into the growing pilus only when they diffuse into a small circular acceptance region (diameter 20 nm, reflecting the diameter of the T4P machine) centered at this point.

The simulation proceeds in discrete time steps (dt = 1 ms). At each step, monomers undergo 2D Brownian motion on the sphere. Incorporation occurs stochastically when a monomer enters the acceptance zone, limited by a characteristic check interval (1/text ≈ 300–333 nm/s reflecting the ATPase-limited velocity under saturating conditions, see experimental results). Extension terminates stochastically at exponentially distributed intervals (median ~3.9 s, corresponding to our experimental findings) to mimic PilB unbinding. The number of incorporated monomers over time is recorded, and the maximum pilus length is computed assuming 1 monomer ≈ 1 nm extension [3].

Simulations were run for varying monomer concentrations N (reflecting different titration levels) and run times T drawn from the exponential distribution (scale 3.9 s) to capture the stochastic nature of extension events. 10,000 replicates per condition were simulated to ensure robust statistics. All code was implemented in Python 3 using freely available packages such as numpy, scipy, matplotlib, pandas, and multiprocessing for efficiency.

### Supplementary Results

#### Mathematical Derivation: Efficiency of Length Regulation vs. Count Regulation in Exponential Pilus Distributions

To quantify the relative efficiency of regulating pilus length versus pilus count for increasing the number of contributing pili, we considered an exponential distribution of pilus lengths with original mean $\mu$ (where median = $\mu \ln(2)$) and functional threshold T (length $\geq$ T to contribute). The baseline fraction of functional pili is $f = \exp(-T / \mu)$. Increasing mean length by factor X (new mean = X $\mu$) yields a new fraction $f\_length = \exp(-T / (X \mu))$. The gain in contributing pili is $f\_length / f = \exp[T / \mu \times (1 - 1/X)]$. Increasing count by X (distribution unchanged) yields a linear gain of X. The efficiency or leverage ratio $R(\mu, T, X) = [f\_length / f] / X = (1/X) \exp[T / \mu \times (1 - 1/X)]$ compares the per-cell gain from length scaling to count scaling; R > 1 indicates length regulation is more efficient.

For example, with μ = 160 nm (20% PilA induction), T = 750 nm (optimized functional threshold), and X ≈ 4.7 (to WT mean ≈750 nm), R ≈ (1/4.7) exp[750 / 160 × (1 − 1/4.7)] ≈ (0.213) exp[4.6875 × 0.787] ≈ 0.213 × 40.3 ≈ 8.6, meaning length scaling by 4.7x yields ~8.6 times more contributing pili than equivalent count scaling. This underscores the exponential advantage/leverage of length tuning, especially when T >> μ, as it disproportionately amplifies the long-tail fraction without the biological costs of massive count increases.

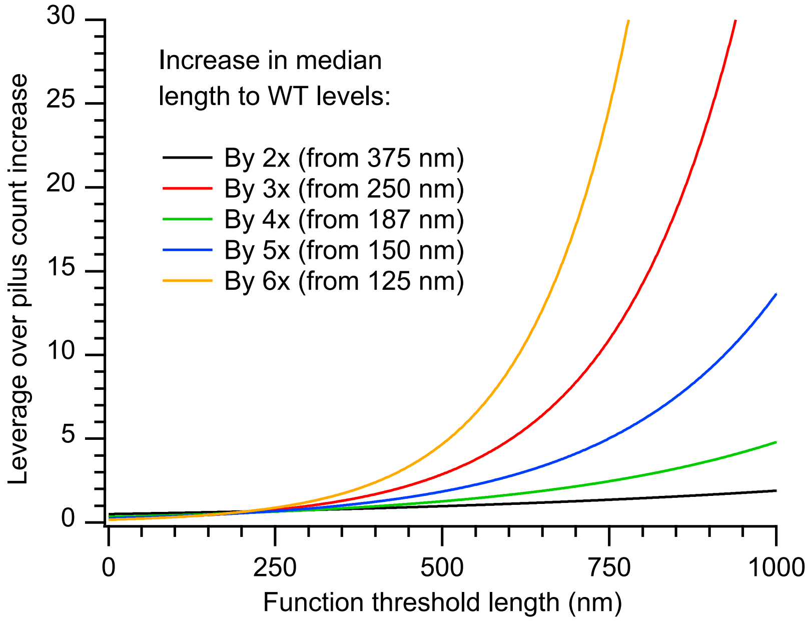

**Supplementary Figure 3. Leverage of tuning the mean pilus length distribution over machine count as a function of the threshold of the length of individual pili that is needed to contribute to biological function and discrete factors X to increase the distribution length to WT levels (~750 nm).**

**Strains used in this study:**

| MK Strain Number | Strains | Strain Description | Reference |
|---|---|---|---|
| MK311 | E. coli | pEXG2 | [4] |
| MK315 | E. coli | pTNS2 | [5] |
| MK420 | S17 | pMK119: pTN7T Pbad::pilA | |
| MK421 | S17 | pMK119: pTN7T Pbad::pilA-A86C | |
| MK373 | S17 | pEXG2 *pilA* | |
| MK367 | S17 | pEXG2 *pilSR* | |
| MK330 | S17 | PaQa mKate2/YFP | [6] |
| MK662 | S17 | pAY1 (PaQa + pilA-YFP) | |
| MK1 | PAO1 | Pseudomonas aeruginosa, WT | [7] |
| MK101 | PAO1 | *pilA -A86C* | [1] |
| MK29 | PAO1 | *pilA*, WT | |
| MK60 | PAO1 | *pilSR*, *pilA* | |
| MK250 | PAO1 | pMK119: pTN7T Pbad::pilA | |
| MK251 | PAO1 | pMK119: pTN7T Pbad::pilA-A86C | |
| MK245 | PAO1 | pMK119: pTN7T Pbad::pilA, *pilA* | |
| MK246 | PAO1 | pMK119: pTN7T Pbad::pilA-A86C, *pilA* | |
| MK255 | PAO1 | pMK119: pTN7T Pbad::pilA, *pilApilSR* | |
| MK804 | PAO1 | pMK119: pTN7T Pbad::pilA-A86C, *pilApilSR* | |
| MK807 | PAO1 | pMK119: pTN7T Pbad::pilA, PaQa mKate2/YFP | |
| MK806 | PAO1 | pMK119: pTN7T Pbad::pilA, *pilA*, PaQa mKate2/YFP | |
| MK808 | PAO1 | pMK119: pTN7T Pbad::pilA, *pilApilSR*, PaQa mKate2/YFP | |
| MK552 | PAO1 | *pilA-A86C*, pAY1 (PaQa + pilA-YFP) | |

**Primers used in this study:**

| Primer | Sequence | |
|---|---|---|
| pEXG2_Ver1 | GTTGCATGGGCATAAAGTTGCC | [1] |
| pEXG2_Ver2 | CGGGTCCTCAACGACAGG | [1] |
| pTn7_Ver1 | GGGTGTAGCGTCGTAAGCTAAT | |
| pTn7_Ver2 | GAAATCAGTCCAGTTATGCTGTG | |
| pilR_P3 | GCACGCATGAGCCGACAAAAACTGGGCATCGACTGAAAGTG | |
| pilR_P4 | GATACAAAGCTTGCATGGTGCTGGCCAAGG | |
| pilS_P1 | GATACAAAGCTTTGACCTTCGGTCTCTTCGAC | |
| pilS_P2 | CTTCCGTCAGCTGAGTTTGCGAGCCGTTCAGCGCGCACGG | |
| PpilA_F1.FOR | GTCGTCCTTGAAAAAGATCGTACGTTCTTGG | |
| PpilA_F1.REV | GGAGAGATTCATGAGCAAAGGTGAAGAACTGTTCAC | |

| | |
|---|---|
| PpilA_F2.FOR | ACAATGAACCCCCGCTGTTGGCGGACCAGCTTTT |
| PpilA_F2.REV | TTTGCTCATGAATCTCTCCGTTGATTATGTATAGGCC |
| PpilA_Ver1 | TGGAGATCGAATTTCAAAGG |
| PpilA_Ver2 | GTATATCGCCCTGCTCCATT |
| pMK119_F1.For | AAGCTAATTCGATCATGCATTTATGACAACTTGACGGCTACATCAT |
| pMK119_F1.Rev | GAATTCCTAGGCATGGTTAATTCCTCCTGTTAGCCCA |
| pMK119_F2.For | CAGGAGGAATTAACCATGCCTAGGCTAGGAATTCAGGAG |
| pMK119_F2.Rev | GATCCACTAGTGAGCTCTTAGTTATCACAACCTTTCGGAGTG |

### Supplimental References:
